# Protein structure from experimental evolution

**DOI:** 10.1101/667790

**Authors:** Michael A Stiffler, Frank J Poelwijk, Kelly Brock, Richard R Stein, Joan Teyra, Sachdev Sidhu, Debora S Marks, Nicholas P Gauthier, Chris Sander

**Affiliations:** cBio Center, Department of Data Sciences, Dana-Farber Cancer Institute, Boston, MA, USA; Department of Cell Biology, Harvard Medical School, Boston, MA, USA; Department of Systems Biology, Harvard Medical School, Boston, MA, USA; Broad Institute, Cambridge, MA, USA; Harvard School of Public Health, Boston, MA, USA; Donnelly Centre, University of Toronto, Canada

**Author notes:** Joint first authors (Michael A Stiffler, Frank J Poelwijk). Joint last authors (Nicholas P Gauthier, Chris Sander). Email to reaches the principal authors (Michael A Stiffler, Frank J Poelwijk, Kelly Brock, Debora S Marks, Nicholas P Gauthier, Chris Sander).

## Abstract

Natural evolution encodes rich information about the structure and function of biomolecules in the genetic record. Previously, statistical analysis of co-variation patterns in natural protein families has enabled the accurate computation of 3D structures. Here, we explored whether similar information can be generated by laboratory evolution, starting from a single gene and performing multiple cycles of mutagenesis and functional selection. We evolved two bacterial antibiotic resistance proteins, β-lactamase PSE1 and acetyltransferase AAC6, and obtained hundreds of thousands of diverse functional sequences. Using evolutionary coupling analysis, we inferred residue interactions in good agreement with contacts in the crystal structures, confirming genetic encoding of structural constraints in the selected sequences. Computational protein folding with contact constraints yielded 3D structures with the same fold as that of natural relatives. Evolution experiments combined with inference of residue interactions from sequence information opens the door to a new experimental method for the determination of protein structures.

## Introduction

By continually generating random DNA sequence variation and selecting for survival, evolution has accumulated a coded record of the physicochemical constraints of the molecular components in evolving organisms. With advances in high-throughput sequencing technology, we now have access to extensive portions of this record in the form of DNA and protein sequence databases. Detecting sequence patterns in homologous proteins has allowed researchers to reconstruct accurate phylogenetic trees and identify functional amino acid residues. A more recent breakthrough uses statistical analysis of co-evolution in protein and RNA families to enable the computation of important interactions between residues or bases, and from those calculate accurate three-dimensional folds and complexes (1, 2).

Here we asked if evolution performed in the laboratory, with its much simplified evolutionary dynamics, similarly encodes information on functional interactions. In contrast, natural evolution is complex, occurring over long time periods, with highly variable population sizes (3), mutation rates (4, 5), and fluctuating environmental conditions (6–11). Each of these factors may or may not be essential for deposition of structural constraints in the evolved sequences. For example, co-evolutionary patterns have been speculated to arise by the continuous degradation and restoration of protein function, i.e. compensatory evolution (12–14), driven by fluctuations in population sizes (15), or periods where functional selection is absent (10). Additionally, natural protein family members may vary in function, operate in various cellular environments, at different optimal temperatures, and importantly, have a broad sequence diversity that is practically unattainable by experimental evolution. The motivation for this work is to use laboratory evolution to elucidate the evolutionary determinants that give rise to co-evolutionary patterns and structural constraints of proteins.

## Results

We subjected two bacterial antibiotic resistance proteins, the Pseudomonas β-lactamase PSE1 and aminoglycoside acetyltransferase AAC6, to repeated rounds of mutation and selection in the laboratory (Fig. 1). To promote sequence divergence, we applied a high mutation rate using an error-prone polymerase chain reaction (epPCR; introducing approximately 3-4% amino acid substitutions per round), and selected for functional proteins under permissive selective conditions (6 μg/mL ampicillin for PSE1 and 10 μg/mL kanamycin for AAC6 — slightly above the minimal inhibitory concentration, MIC, for E. coli lacking a resistance gene). These conditions generally resulted in survival of about 1% of the initial population post-selection (approx. 5×10^4^ cells for PSE1 and 2×10^5^ for AAC6) in each round. Successive rounds of mutation and selection were applied by using the selected sequences in one round as the template for mutations in the next round. We deep-sequenced the selected populations, obtaining 10^4^-10^6^ high-quality unique reads, at rounds 10 and 20 for PSE1 and rounds 2, 4, and 8 for AAC6.

**Figure 1.**
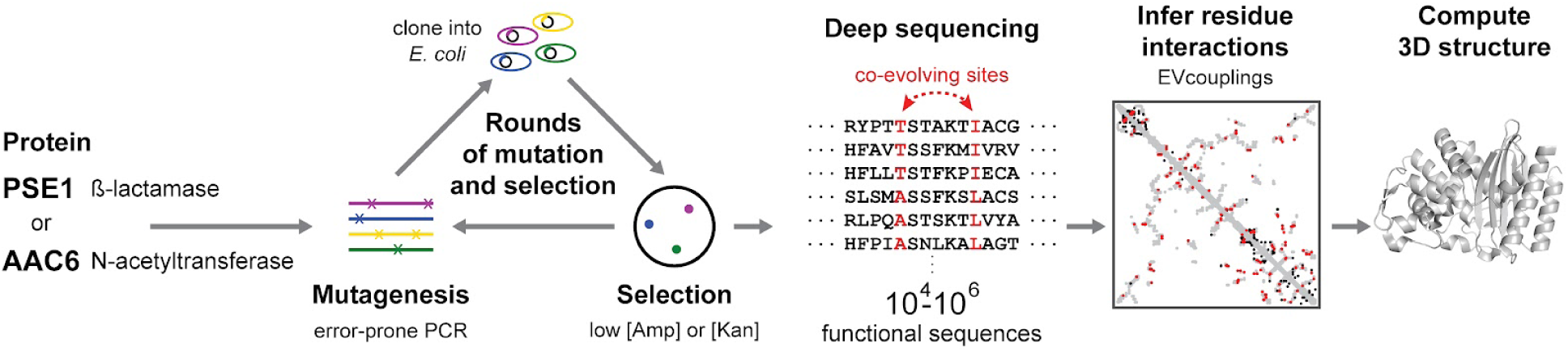
Approach: from laboratory evolution experiments to residue interactions and three-dimensional structures. The experiments involve repeated rounds of mutation and selection, starting from a single sequence (β-lactamase PSE1, 266 residues, or aminoglycoside acetyltransferase AAC6, 148 residues). In each round, mutations are generated by error-prone PCR, followed by cloning into bacteria (E. coli), and selection for functional variants at relatively low antibiotic concentration (6 μg/ml ampicillin [Amp] for PSE1 and 10 μg/ml kanamycin [Kan] for AAC6). A large number of full-length sequences at various rounds are obtained by deep sequencing after selection; at rounds 10 and 20 for PSE1, and rounds 2, 4 and 8 for AAC6. Residue interactions are inferred from co-evolution patterns in the selected sequences using the evolutionary couplings (EVcouplings (1)) maximum entropy model, which are then used as distance constraints to compute three-dimensional structures using distance geometry and simulated annealing molecular dynamics (16).

Sequencing revealed that the mutation count relative to the ancestral sequence increases with the number of rounds of mutation and selection (Fig. 2A). In the final round, the evolved sequences have an average of 34.2 mutations (12.9% of sequence length) in PSE1 and 8.7 mutations (5.9%) in AAC6. Thus, of the 3-4% amino acid mutations introduced per round in each sequence, the functionally-selected sequences end up with an average of 1.7 (0.6%) and 1.2 (0.8%) amino acid mutations for PSE1 and AAC6, respectively. In the later rounds there was a trend towards fewer tolerated mutations, e.g. for PSE1, 1.9 mutations were added per round up to round 10 and 1.5 mutations were added per round between rounds 10 and 20.

**Figure 2.**
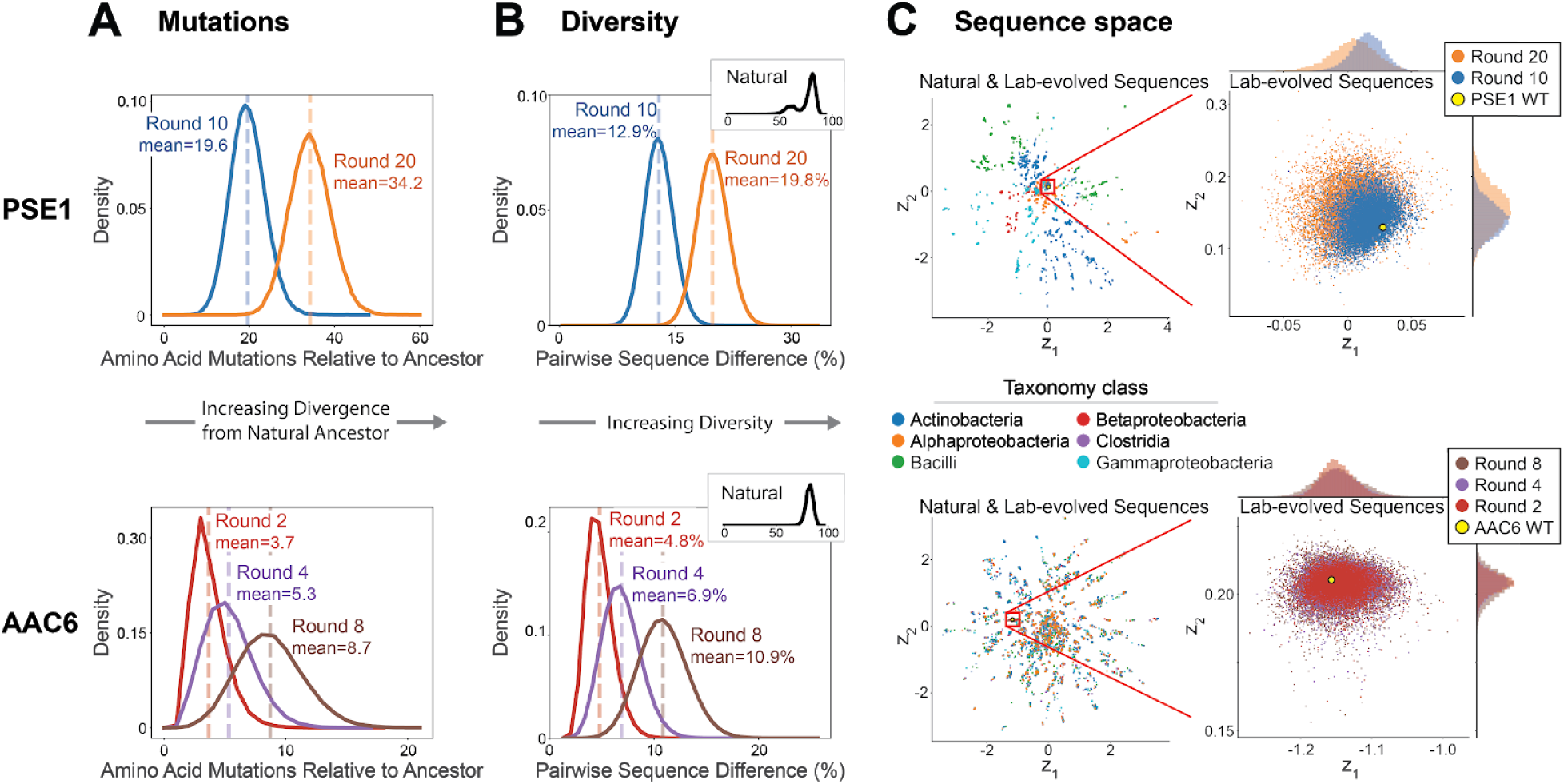
Divergence, diversity and sequence space of lab-evolved sequences. (**A**) Distributions of amino acid mutations relative to starting (ancestral) sequence, per unique sequence obtained from laboratory evolution of PSE1 (top) and AAC6 (bottom). (**B**) Distributions of pairwise sequence diversity (percent positions with non-identical amino acids) calculated for 25×10^6^ randomly chosen pairs of unique sequences for PSE1 (top) and AAC6 (bottom). Sequence diversity among natural homologs is substantially larger (inset at top right). (**C**) Two-dimensional representation of sequence sets (each point is one protein sequence) in sequence space projected down from the N-dimensional space using a variational autoencoder (17) with two latent variables (z_1_ and z_2_). Sequences of natural homologs (current databases, colored by taxonomy) occupy a much larger space than those from our laboratory evolution experiments. The lab-evolved sequences increasingly separate from the ancestral sequence with increasing rounds of mutagenesis (point cloud on right).

The mutational distance to the ancestor is not by itself a measure of diversity as the libraries could consist of sets of very similar sequences. To assess diversity, we monitored the all-against-all pairwise sequence differences in each population. We observed an average of 19.8% and 10.9% pairwise sequence differences in the final round of PSE1 and AAC6 evolution, respectively (Fig. 2B). For both proteins, this equates to an increase in pairwise sequence difference of ∼1.2% per round — close to the maximum possible increase if the populations were freely expanding in sequence space. We conclude that our approach effectively generates and preserves a high level of sequence diversity. In contrast, the mean pairwise sequence difference within the set of known natural homologs of PSE1 or AAC6 is around 80% (Fig. 2B). Projected onto two-dimensional sequence space, the laboratory-evolved sequences increasingly disperse with increasing rounds of mutation and selection, but otherwise occupy a small and dense area relative to natural sequences (Fig. 2C).

Information about evolutionary constraints in iso-functional sequences increases both with sequence diversity and with the total number of non-identical sequences (1, 18). Although the level of diversity and positional entropy in the lab-evolved sequence sets is lower than in families of natural homologs (Fig. 2C), we have generated many more lab-evolved sequences than are currently available for natural homologs (final laboratory evolution rounds have 1.6×10^5^ unique functional sequences for PSE1 and 1.3×10^6^ for AAC6; the PFAM database has 3.7×10^4^ homologs for PSE1 (PFAM ID = PF00144) and ∼1.2×10^5^ unique functional for AAC6 (PF00583).

To quantify the extent to which the laboratory evolution process has encoded co-variation patterns that are informative of interactions between pairs of residue positions, we used a global probability model (EVcouplings) that has been successful in detecting such patterns in natural sequences (Fig. 1) (1, 19, 20). We compared the inferred interactions to actual contacts in published crystal structures closest in sequence to each of the two ancestral proteins (Protein Data Bank 1G68 for PSE1, and 4EVY for AAC6). Comparison with crystal structures tests whether functional selection (i.e., enzymatic deactivation of antibiotic) conserves three-dimensional structure - it is well established from analysis of natural sequences and structures that there is a high degree of structural conservation among iso-functional homologs even with highly diverged sequences (21). However, in lab evolution one may expect more structural variability compared to natural evolution, as some protein structural properties, such as those related to aggregation or thermodynamic stability (12, 22), may be under weaker selection in the laboratory than in nature, given the much shorter timescales, smaller population sizes, and homogeneous environments (23).

In practice, we defined contact agreement as the percentage of top-ranked inferred interactions (typically L/2 inferred interactions where L is the length of the protein) that are also contacts in the crystal structure. Agreement increases with successive rounds of mutation and selection (Fig. 3A) as sequence diversity increases. For PSE1, the agreement increased from 34% in round 10 to 49% in round 20 (for a subset of 4×10^4^ unique mutant sequences from each round). For AAC6 the agreement was 22% in round 2, 30% in round 4, and 42% in round 8 (for an equal size subset of 10^5^ unique sequences). These percentages are well above the random expectation of 1.9% and 4.1% for PSE1 and AAC6 (approximated by the ratio of crystal structure contacts over the total number of pairs), even at early rounds (see AAC6 round 2, Fig 3A). Additional sequencing also led to a higher agreement between inferred interactions and contacts: for the last rounds we obtained 1.6×10^5^ unique functional sequences for PSE1 and 1.3×10^6^ for AAC6, leading to a contact agreement of 54% for PSE1 and 51% for AAC6 (Fig. 3). These results indicate that simplified evolutionary dynamics in the laboratory do generate functional sequences with co-evolutionary patterns that reflect constraints imposed by three-dimensional protein structure.

**Figure 3.**
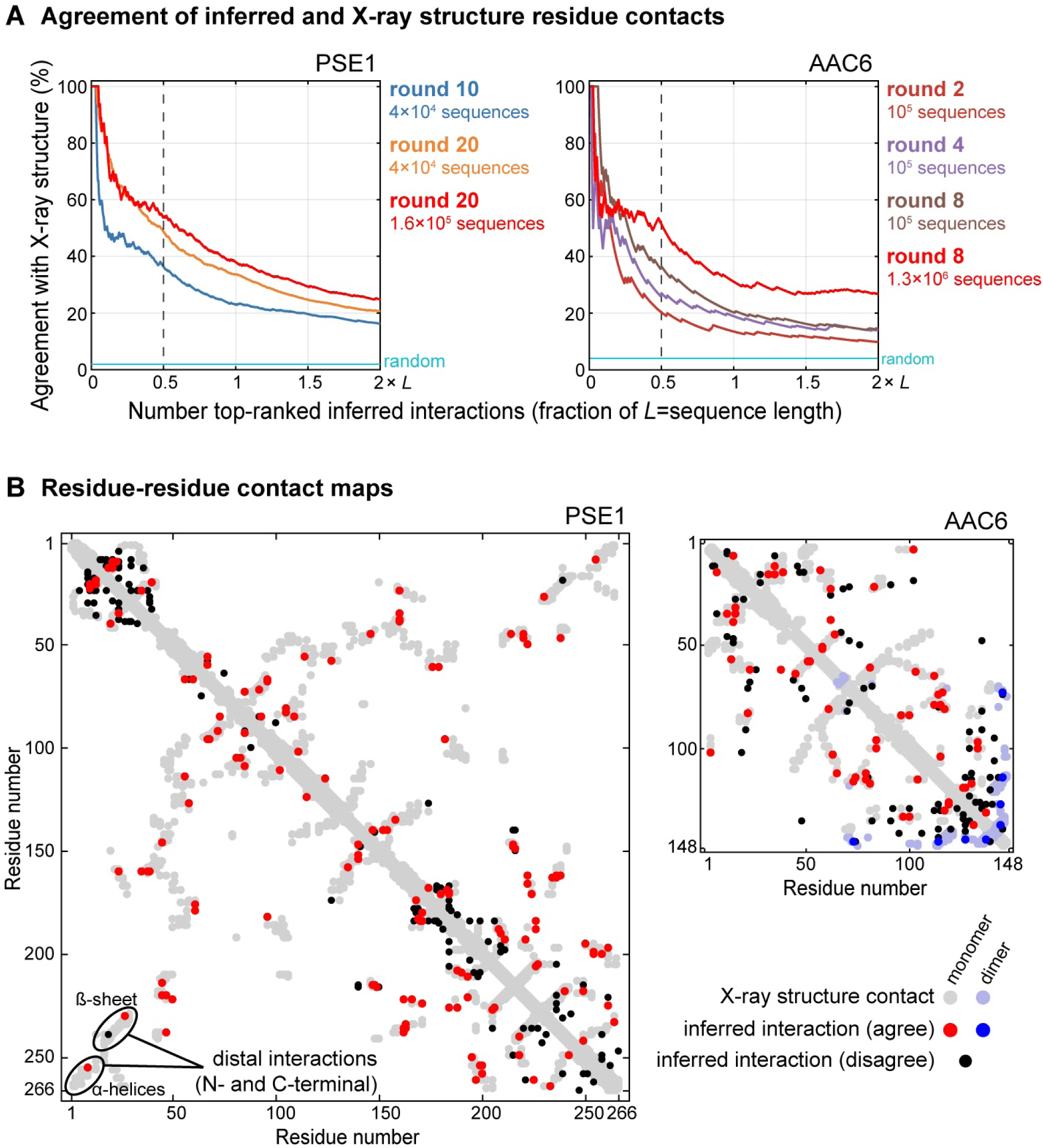
Agreement between residue contacts inferred from laboratory evolution and contacts in crystal structures. (**A**) Agreement versus number of inferred interactions (as fraction of sequence length, L) during lab evolution of PSE1 (left) and AAC6 (right). PSE1 results evaluated for an equal number (4×10^4^) of unique sequences from rounds 10 and 20 to illustrate change in agreement with increased rounds of mutation and selection, and all (1.5×10^5^) unique sequences at round 20 to illustrate change with increased number of sequences. AAC6 similarly assessed for an equal number (10^5^) of unique sequences at rounds 2, 4 and 8, and all (1.3×10^6^) unique sequences at round 8. Random is the average result obtained with randomly chosen residue pairs. (**B**) Inferred interactions from PSE1 evolution at round 20 (left) and AAC6 evolution at round 8 (right), overlaid on contact maps of crystal structures. Inferred interactions either agree with monomer (red) or dimer (blue) contacts in the crystal structure (gray or light blue, respectively), or disagree (black). For PSE1, the distal residue interactions between the N- and C-terminal α-helices and β-strands (lower left) are particularly crucial constraints for the correct 3D fold via reduction of chain entropy. Dashed line in (A) and results in (B) are at L/2 inferred interactions; agreement of >50% at L/2 often suffices to compute 3D structures (2). In (A) and (B), X-ray structure contacts are defined as residue pairs < 5 Å minimum side chain atom distance; inferred interactions are limited to a primary sequence distance > 5 residues.

The residue interactions inferred from laboratory evolution include important structural features of both proteins (Fig. 3B). The β-lactamase fold, for example, consists of two structural domains, an all-α-helical domain and a mixed α/β domain (24, 25). The polypeptide chain is interwoven between the two domains leading to extremely sequence-distal contacts between the N- and C-terminal α-helices, and the N- and C-terminal β-strands. Consistent with the crystal structures, we likewise observe strongly co-evolving interactions between these sequence-distal structural elements in the PSE1 results (Fig. 3B). Proteins from the same subfamily of aminoglycoside acetyltransferases as AAC6 exist as homodimers, with the C-terminal β-strand of one protein chain inserted between two strands of the other chain (26). We also discern these dimer interactions among the top-ranked inferred residue-residue contacts (Fig. 3B). Similar to previous work on natural sequences (27), identification of these inter-protein contacts demonstrates that laboratory evolution can also be informative of protein-protein interactions.

We next asked whether the inferred contacts from laboratory evolution are sufficient to compute the three-dimensional structure. For natural protein families, inferred residue interactions in the range of agreement with crystal structures of 50-60% are typically sufficient to compute three-dimensional folds that agree with those observed by crystallography or NMR (1, 28). To assess whether laboratory evolution provides a similar level of information, we computed sets of structures using inferred residue interactions as distance constraints in molecular dynamics with simulated annealing (1, 16, 28). The constraints are updated using a new filtering algorithm. For AAC6 we folded only the monomeric unit consisting of residues 1-134, as residues 135-148 are directly involved in dimer contacts and computation of a dimeric molecule from a compounded contact map is beyond the scope of this work. Algorithmic filtering of the constraints led to improved agreement between inferred interactions and crystal structure contacts, from 54% to 65% for PSE1, and from 45% to 59% for AAC6 (Fig. 4A).

**Figure 4:**
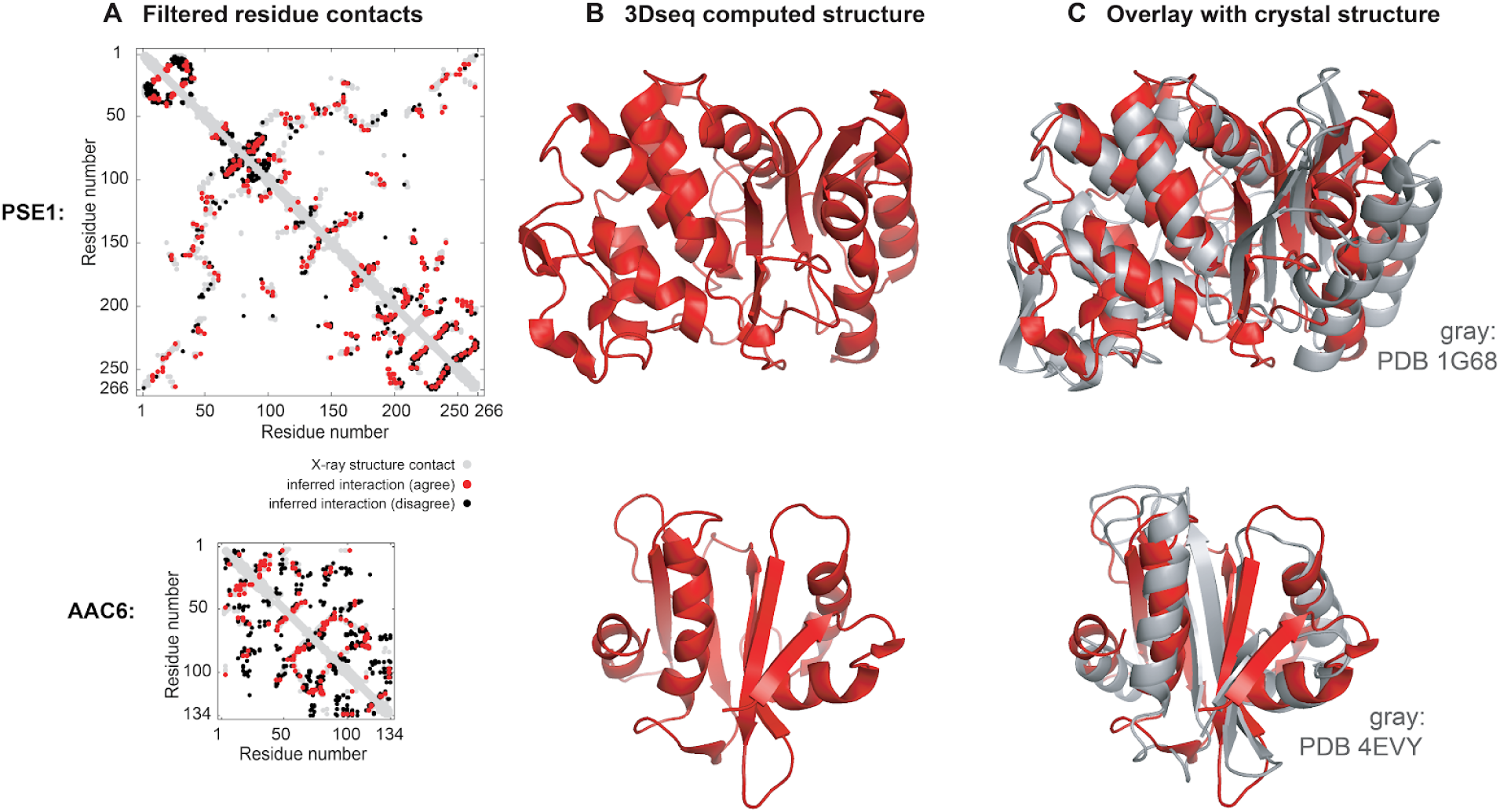
3D structures computed from experimental evolution compared to those from X-ray crystallography. (**A**) Inferred interactions after applying a constraint filtering algorithm; red agree with contacts in the crystal structure, black disagree. Shown are results for 2×L inferred interactions. (**B**) 3D structures computed using the filtered inferred interactions as distance constraints. Red ribbons are structures with the lowest Cα positional RMSD over more than 90% of residues for either protein. (**C**) Computed structures compared to crystal structures (gray ribbons): for PSE1 (PDB 1G68, Cα positional RMSD 4.5 Å over 240/266 residues, TM-score = 0.65); for AAC6 (structural homolog of AAC6 PDB 4EVY, Cα positional RMSD 3.8 Å over 122/130 residues in 4EVY, TM score = 0.59). For AAC6, the C-terminal β-strand known to be involved in dimer contact is excluded; we did not attempt to compute dimer structures. Overall, structures computed from interactions inferred from laboratory-evolved sequences have the same general fold as crystal structure, with template modeling score (TM-score <u>(29)</u>) of >0.5 for 72% of PSE1 models (690 total models) and for 63% of AAC6 models (720 total models).

The final set of structures, computed from the inferred interactions after filtering, was assessed for agreement with the known crystal structure of the most sequence-similar homolog. Of the set of computed structures, 72% of PSE1 and 63% of AAC6 generated structures (of 690 models generated for PSE1 and 720 for AAC6) have a template modeling score (TM-score) of 0.5 or greater; TM-scores in excess of 0.5 are generally considered to indicate overall fold similarity (29–31). The structures with the lowest C positional root-mean-square deviation (RMSD) over more than 90% of residues and which do not contain knots (32, 33) have 4.5 Å RMSD for PSE1 (240/266 residues) and 3.8 Å RMSD for AAC6 (122/130 residues of 4EVY) (Fig. 4). Overall, we conclude that the laboratory evolution process reflects natural evolution in constraining residue interactions that conserve the same three-dimensional fold.

## Discussion

The experimental evolution approach taken here opens the door to a new ab initio experimental method of determining protein structure, which we call 3Dseq. Using laboratory evolution for structure determination is complementary to established methods such as X-ray crystallography, NMR and cryo-electron microscopy, in several respects, with several advantages and disadvantages. For example proteins can be interrogated in a native cellular environment, not requiring biochemical purification of proteins; inter-molecular interactions can be elucidated without purification and/or crystallization of complexes (dimer contacts are already inferred in AAC6) by experimental evolution using, e.g., two-hybrid (34–36), or phage- or yeast- display approaches (37, 38); and by targeted mutagenesis and/or controlling selective conditions, one can infer which constrained interactions are of functional importance under a given selection condition. However, as 3Dseq infers distance constraints from co-variation data in entire sets of diverse sequences, its precision of atomic positions, even when the fold is correct, is significantly less than that of single-sequence, single-conformation crystallography. The precision of determining single-sequence structures would increase with improvements in constrained molecular dynamics. To compute single sequence structures for all sequences in the lab-selected libraries, one would have to execute many thousands constrained molecular dynamics runs, which is beyond the scope of this work.

Future generalization of 3Dseq experimental technology would benefit from assays that directly select or screen for protein structural integrity, which is more generally applicable than dependence on selection for a particular cellular function (39–41), (e.g., antibiotic resistance). A major efficiency gain would come from automated evolution systems, which combine mutation and selection in single cells and rely on proliferative advantage in pooled experiments (42–44).

In using experimental evolution to elucidate constraints one is agnostic as to exactly how evolutionary constraints are encoded in the protein sequences. Recent work (45, 46) showed that two-way epistasis between amino acid mutations derived from deep mutational scans can capture structural constraints sufficient for the computation of 3D structures, at least for small proteins. Given the ability to control sequence diversity, depth of sequencing and quantitation of fitness in laboratory evolution, future evolution experiments will provide an opportunity to unravel evolutionary pathways, the level of cooperativity in sets of mutations, and to discover constraints essential for the maintenance of structure and function.

In contrast to natural evolution, laboratory evolution is typically performed in less diverse environments, over much shorter time scales, and with simpler population dynamics (7, 23). Nonetheless, our results indicate that laboratory evolution consisting of repeated cycles of random mutation and uniform selection can generate large and diverse sets of sequences with rich co-evolutionary interaction patterns. Experimental evolution approaches of this type can serve as a contribution to better understanding the complexities of natural evolution and to developing quantitative models of both retrospective and prospective evolutionary dynamics.

## Acknowledgements

We thank our colleagues for help. EVcouplings pipeline: Thomas Hopf, Christian Dallago, Benjamin Schubert; DeepSequence software package: John Ingraham, Adam Riesselman; Sequencing: the Molecular Biology Core Facilities at Dana-Farber (Zack Herbert, Maura Berkeley), MIT BioMicro Center, and the Genomics Core Facility at the Icahn Institute, Mt. Sinai (Melissa Smith, Diane Castillo, Gintaras Deikus); Pacbio process: Michael Weiand from Pacific Biosciences; Discussion: members of the Sander and Marks labs.

